# Distribution and origins of *Mycobacterium tuberculosis* L4 in Southeast Asia

**DOI:** 10.1101/2022.08.01.502309

**Authors:** Philip M. Ashton, Jaeyoon Cha, Catherine Anscombe, Nguyen T.T. Thuong, Guy E. Thwaites, Timothy M. Walker

## Abstract

Molecular and genomic studies have revealed that *Mycobacterium tuberculosis* Lineage 4 (L4, Euro-American lineage) emerged in Europe before becoming distributed around the globe by trade routes, colonial migration, and other historical connections. Although L4 accounts for tens or hundreds of thousands of TB cases in multiple Southeast Asian countries, phylogeographic studies have either focussed on a single country or just included Southeast Asia as part of a global analysis. Therefore, we interrogated public genomic data to investigate the historical patterns underlying the distribution of L4 in Southeast Asia and surrounding countries. We downloaded 6037 genomes associated with 23 published studies, focusing on global analyses of L4 and Asian studies of *M. tuberculosis. We* identified 2256 L4 genomes including 968 from Asia. We show that 81% of L4 in Thailand, 51% of L4 in Vietnam and 9% of L4 in Indonesia belong to sub-clades of L4 that are rarely seen outside East and Southeast Asia. These sub-clades have been transmitted between East and Southeast Asian countries, with no recent European ancestor. Although there is considerable uncertainty about the exact direction and order of intra-Asian transmissions, due to differing sampling frames between countries, our analysis suggests that China may be the intermediate between Europe and Southeast Asia for two of the three predominantly East and Southeast Asian L4 sub-lineages. This new perspective on L4 in Southeast Asia raises the possibility of investigating host population specific evolution and highlights the need for more structured sampling from Southeast Asian countries to provide more certainty of the historical and current routes of transmission.

## Intro

*Mycobacterium tuberculosis* caused 5.8 million reported cases of tuberculosis and 1.5 million reported deaths in 2020 (1). The significant disruption to TB services from the COVID-19 pandemic mean the true numbers are much higher (1,2). Molecular and genomic studies have revealed that there are at least eight lineages of *M. tuberculosis,* which display variable degrees of phylogeographical signal (3–5).

*M. tuberculosis* Lineage 4 (L4) is globally distributed (6), and is thought to have originated in Europe (7–9). Colonialism and long-distance trade have been proposed to be important for the spread of L4 to the Americas, Africa, Asia and Oceania (9,10). While Lineage 2 (Beijing lineage) is the most prevalent lineage in most countries in East and Southeast Asia, the high TB burden in these countries means that the absolute number of TB cases caused by L4 isolates is nevertheless large. Much of our understanding of TB molecular epidemiology in the region is derived from spoligotyping. This showed that Indonesia had the highest proportion of L4 in Southeast Asia, with publications focussing on different parts of the country reporting 28-47% L4 (6,11–14). Analysis of the spoligotype of 16,621 isolates from all 32 provinces of China found that L4 accounts for around 17% of M. *tuberculosis there* (15). Lower prevalence of L4 is seen across Vietnam (6.4-12.2% (6,16,17)), Myanmar (8% (18)), Thailand (10% (6)) and Malaysia (9.6-13.5% (6,19)). Reports from Cambodia and Philippines show lower rates still (0-1% (20,21) and 1% (22) respectively).

Studies making use of whole genome sequencing (WGS) have recently improved our understanding of L4 in East and Southeast Asia. A well-structured sampling of 279 L4 genomes from China revealed that 97% of L4 in China belongs to one of three L4 sublineages (L4.2.2, L4.4.2 and L4.5) which were introduced from Europe between the 11^th^ and 13^th^ Centuries, likely mediated by the intense trade connections at this historical period, exemplified by the Maritime Silk Road (15). From Southeast Asia, only Vietnam, Indonesia, Thailand and the Philippines have more than 10 published L4 genomes (23–26). The L4 genomes from Vietnam came from a study of the genomic epidemiology of *M. tuberculosis* in Ho Chi Minh City, and a study of drug resistant *M. tuberculosis* in Hanoi (25,27). The currently available L4 genomes from Indonesia were isolated in the city of Bandung on the island of Java as part of a study examining differences between *M. tuberculosis* causing pulmonary tuberculosis and tuberculosis meningitis (24). L4 genomes from Thailand came from studies comparing pulmonary tuberculosis and tuberculosis meningitis and a cohort study (23,28). The analyses from Indonesia and Thailand did not include any international genomes for context, so their phylogenetic relation to the broader diversity of L4 is unknown (23,24).

However, our understanding of L4 in East and Southeast Asia remains piecemeal as no studies to date have combined all published datasets for analysis. Therefore, to investigate the historical patterns underlying the present distribution of L4 in Southeast Asia, we carried out a combined analysis of published *M. tuberculosis* L4 genomes from East and Southeast Asia, along with contextual genomes from global data sets.

## Results

We identified 6037 read-sets associated with 23 publications on *M. tuberculosis* genomics (8,9,15,23–42) and downloaded them from the European Nucleotide Archive (ENA). We identified 2257 L4 read-sets that were not mixed and that had associated geographical information (Table 1, Supplementary Table 2). Of these 2257 read-sets included in this analysis-sets, 968 (43%) were from Asia, 581 (26%) were from Europe, 501 (22%) were from South America, 144 (6%) were from Africa, 58 (3%) were from North America and 5 (0.2%) were from Oceania (Supplementary Table 1, Supplementary Table 2: A full line list of all lineage 4 readsets included in this analysis Supplementary Table 3). We identified 308 read-sets as belonging to the L4.3.3 sub-lineage, 268 as L4.5, 255 as L4.1.2.1, 237 as L4.8, 156 as L4.3.4.2, 140 as L4.4.2, 110 as L4.4.2 and 784 belonging to other L4 sub-lineages (Supplementary Table 2: A full line list of all lineage 4 readsets included in this analysis Supplementary Table 3). As a note on terminology, we use the term “sub-lineage” to refer to a group of L4 M. tuberculosis with a common Coll et al., designation e.g., L4.4.2 (43), and “sub-clade” to refer to a monophyletic part of a sub-lineage. A sub-clade may have different geographical distribution than the overall sub-lineage.

**Table 1:**
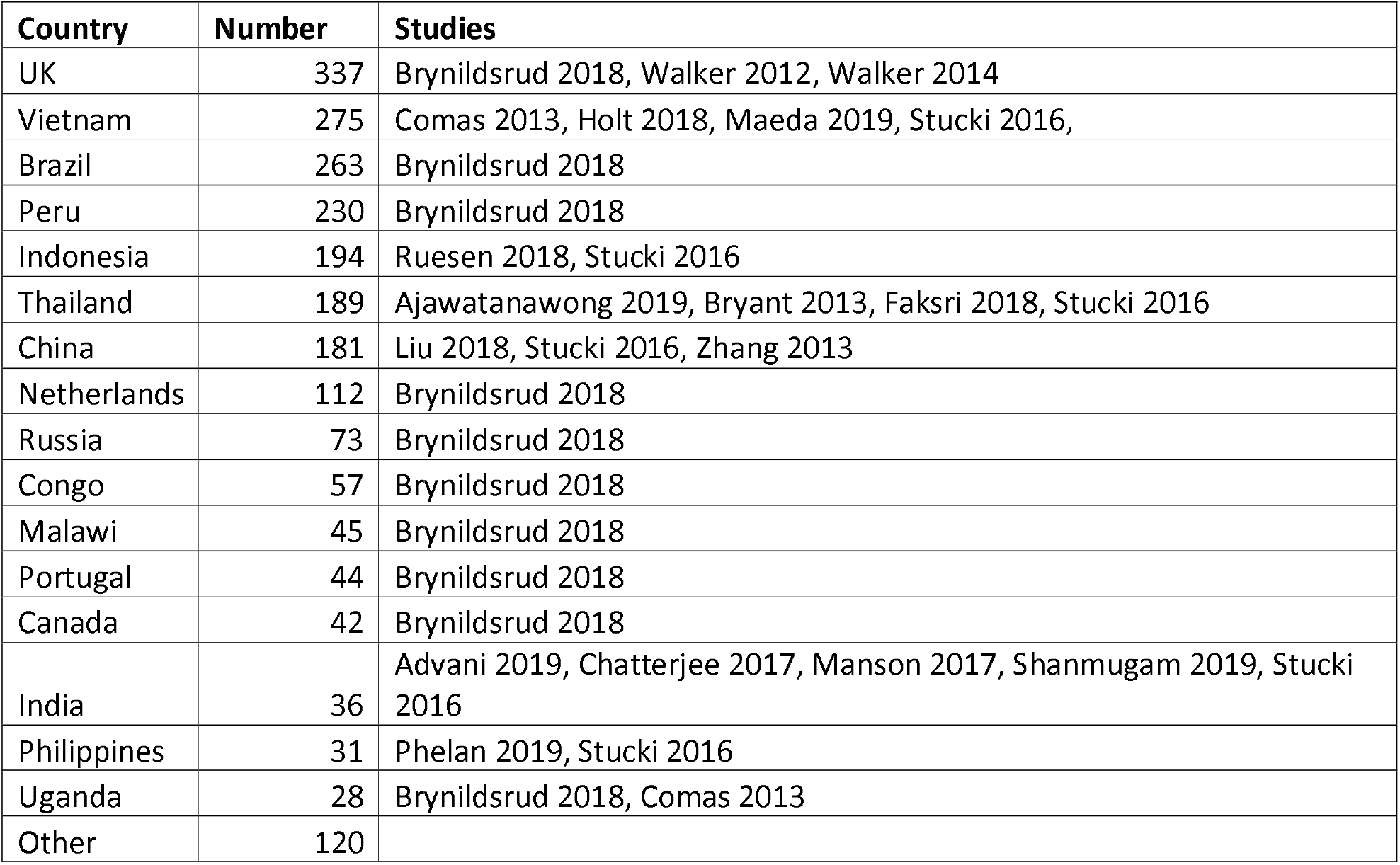
The number of genomes from each country included in this study, and the studies they were first reported in.

*We* constructed a maximum likelihood phylogenetic tree of 2258 L4 genomes (2257 read sets and the H37Rv reference genome, Figure 1) and used it as the input for TreeBreaker (44) analysis (see methods). This identified 40 sub-clades with notable changes in the proportion of tips coming from East and Southeast Asian countries (see methods). These sub-clades were analysed alongside SIMMAP reconstruction of the geographical location of ancestral nodes in the phylogeny to improve our understanding of the current and historical distribution of *M. tuberculosis* L4 in Southeast Asia. Internal branches identified by both TreeBreaker and SIMMAP as transmissions between our geographical areas of interest are highlighted in Supplementary Figure 1.

**Figure 1:**
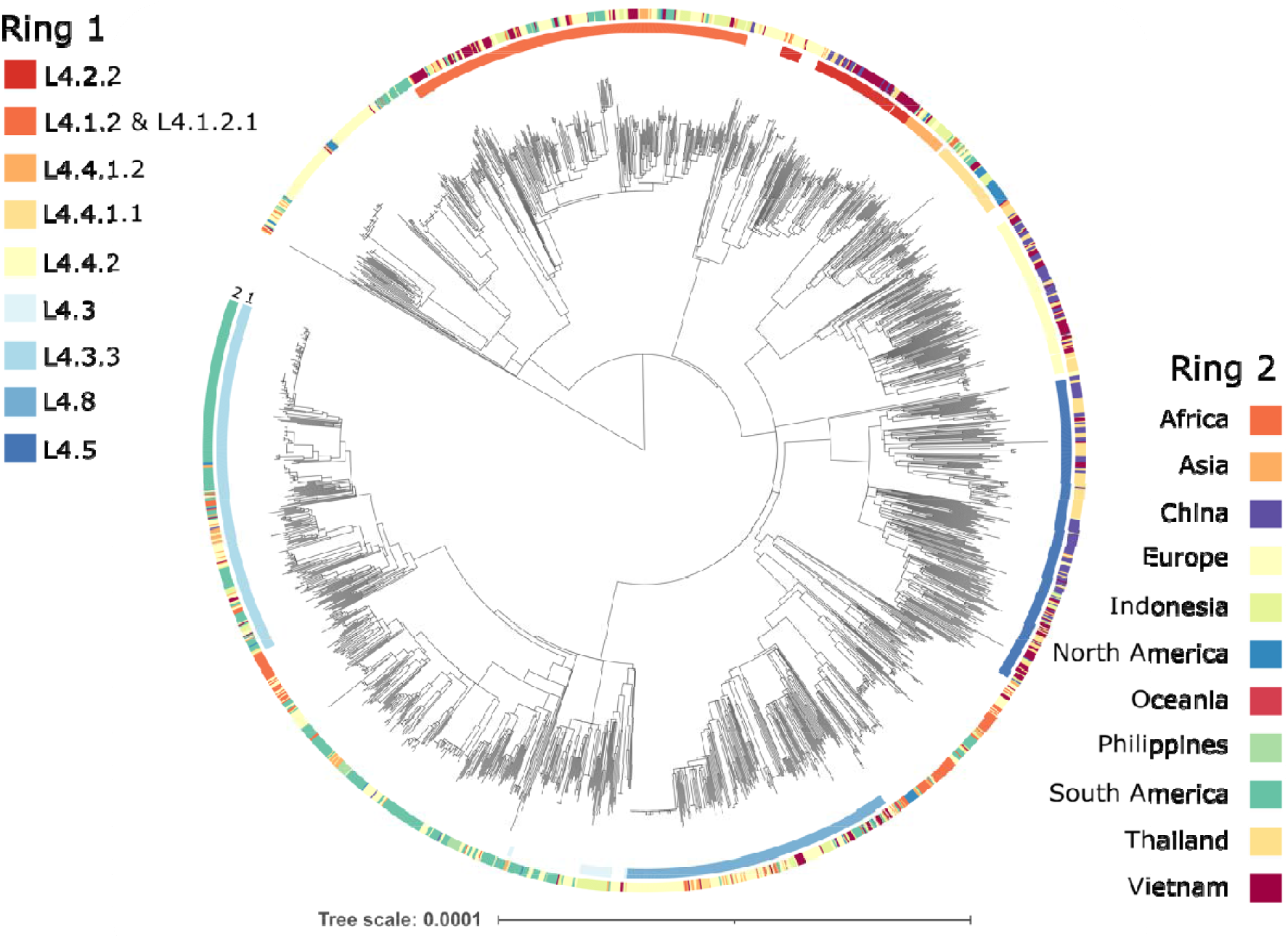
A maximum likelihood phylogenetic tree of 2258 M. tuberculosis L4 genomes (the 2257 read sets identified and the H37Rv reference genome). Ring 1 (inner ring) indicates the Coll et al., 2014 sub-lineage. For clarity, we only highlighted sub-lineages that are discussed further in this paper. Ring 2 (outer ring) indicates the country of origin.

We identified three main kinds of sub-clade in our analysis: i) Southeast Asian sub-clades in East/Southeast Asian sub-lineages, ii) Southeast Asian sub-clades within global lineages, and iii) reversions, which are Southeast non-Asian sub-clades nested within an East/Southeast Asian sub-clade (Figure 2). We define East/Southeast Asian sub-lineages as those where >=75% of the sub-lineage in our collection was from East or Southeash Asia, and where there were more than 50 genomes from that sub-lineage in our analysis. East/Southeast Asian sub-clades are those identified by TreeBreaker as being associated with our countries of interest.

**Figure 2:**
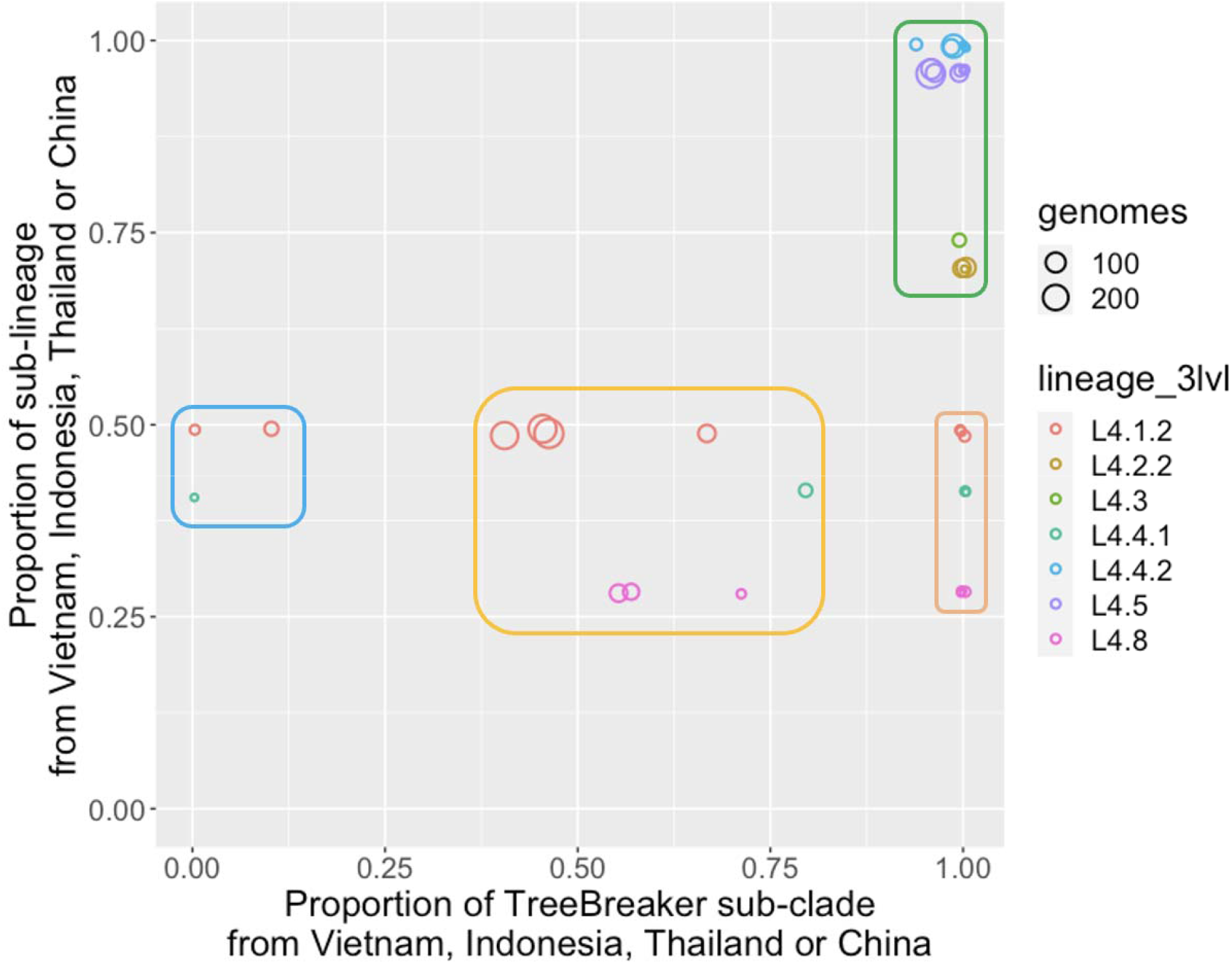
We identified four kinds of L4 sub-clades among those identified by TreeBreaker that varied by the proportion of the sub-clade that was present in Vietnam, Indonesia, Thailand or China, and by the proportion of the sub-lineage that were from those countries. Each circle represents a sub-clade descending from an internal branch that TreeBreaker identified as representing a change between two geographical locations. The rectangles with curved corners represent the kinds of sub-clade identified. The size of the circle represents the number of genomes in the sub-clade. The circles are not necessarily independent as sub-clades can be nested within higher sub-clades. The position on the x-axis is the proportion of the sub-clade identified by TreeBreaker that came from Vietnam, Thailand, Indonesia, or China. The position on the y-axis is the proportion of the sub-lineage to which the sub-clade represented by that circle belongs, that comes from Vietnam, Indonesia, Thailand, or China. The colour of each circle represents the sub-lineage it belongs to. The first “type” of sub-clade we identified is Asian sub-clades in Asian sub-lineages, located in the top right of the graph (inside a green rectangle) because a high proportion of this sub-clade is from Asia, and also the majority of isolates the sub-clades we identified in our TreeBreaker analysis come from Asia. The second type of sub-clade we identified was Asian sub-clades within global lineages, located in the centre of the x-axis and below the 0.5 mark on the y-axis (light orange rectangle) because they represent sub-clades enriched in Asian isolates within sub-lineages that are generally global. Derived from this type of sub-clade are smaller sub-clades of the same sub-lineage that are below 0.5 on the y-axis and on the extreme right of the x-axis (dark orange rectangle), which are small sub-clades entirely from Vietnam, Indonesia, Thailand or China, within the global lineages. Finally, there are “reversions” of sub-clades that are within Asian enriched sub-clades but that are majority non-Asian, placed in the bottom left of the figure (blue rectangle). These represent transmissions of L4 sub-clades from Asia to other parts of the world.

### East/Southeast Asian sub-lineages

In China, 94% of L4 belonged to the East/Southeast Asian sub-lineages, in Thailand it was 81%, in Vietnam it was 51% and in Indonesia it was 9%.

#### L4.2.2

L4.2.2, one of the East/Southeast Asian sub-lineages, had a sub-clade of 88 genomes identified by TreeBreaker, of which 54 were from Vietnam, 21 were from China, 12 were from Thailand and 1 was from Indonesia (Figure 3). SIMMAP analysis showed that the probable geographic location of the Most Recent Common Ancestor (MRCA) of this sub-clade was in China (99% probability), while the parent node of the MRCA was assigned to Europe (99% probability). This relationship between the MRCA node and the parent node of the MRCA indicates that, according to the SIMMAP analysis, a transmission event from Europe to China occurred at some point along the branch between those two nodes. There was a further sub-clade of 64 genomes, including 51 of the 54 Vietnamese genomes, for which the MRCA was probably located in Vietnam (81%), while the parent node of the MRCA was placed in China (99%). One hundred percent of the sub-sampled replicates identified China as the location of the MRCA of L4.2.2.

**Figure 3:**
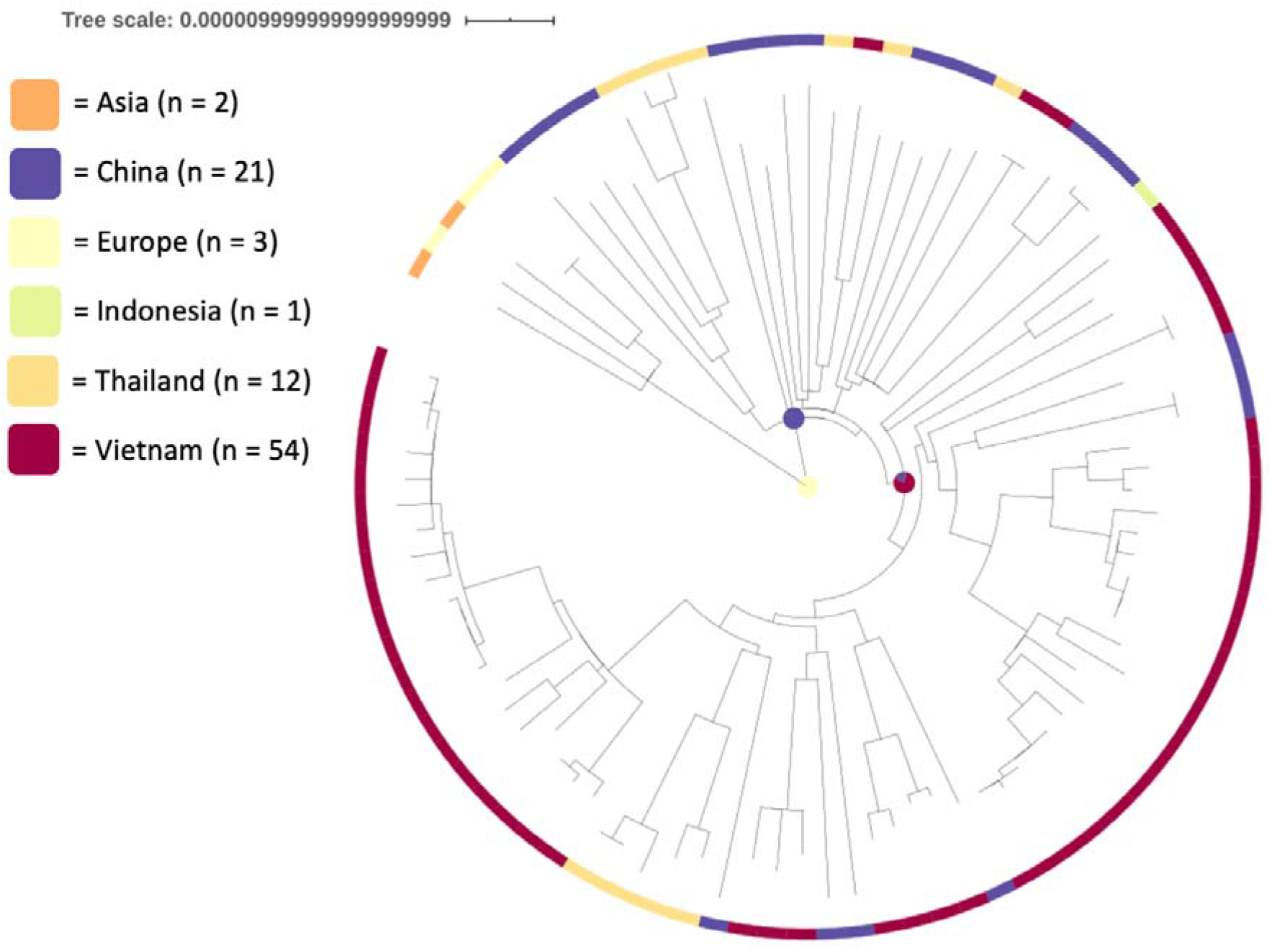
A maximum likelihood phylogeny of 88 L4.2.2 genomes. SIMMAP analysis indicates that there was a transmission from Europe to China, and subsequently from China to Vietnam. Pie charts on internal nodes represent the probability of the geographical location of that hypothetical ancestor.

#### L4.4.2

L4.4.2 was another East/Southeast Asian sub-lineage, with 152 genomes in a sub-clade identified by TreeBreaker (including 12 basally branching L4.4 genomes), of which 53 were from China, 49 were from Thailand, 43 from Vietnam, 5 from Indonesia, 1 from Malaysia and 1 from the UK (Figure 4). SIMMAP produced an ambiguous result for the location of the MRCA of this sub-clade, and of the parent node of the MRCA. The MRCA was in either China (54%) or Vietnam (34%), while the parent node of the MRCA was either in Europe (55%) or Vietnam (29%). However, a sub-clade of 135 of the 152 genomes had an MRCA which was assigned to China with a high probability (>99%). Within this, there was a clade of 52 genomes for which the MRCA was in either Vietnam (75%) or China (25%), while the parent of the MRCA was confidently assigned to China (>99%). In the sub-sampling analysis, 56.8% of the replicates placed the MRCA in Thailand, 43% in Vietnam, and 0.2% in China.

**Figure 4:**
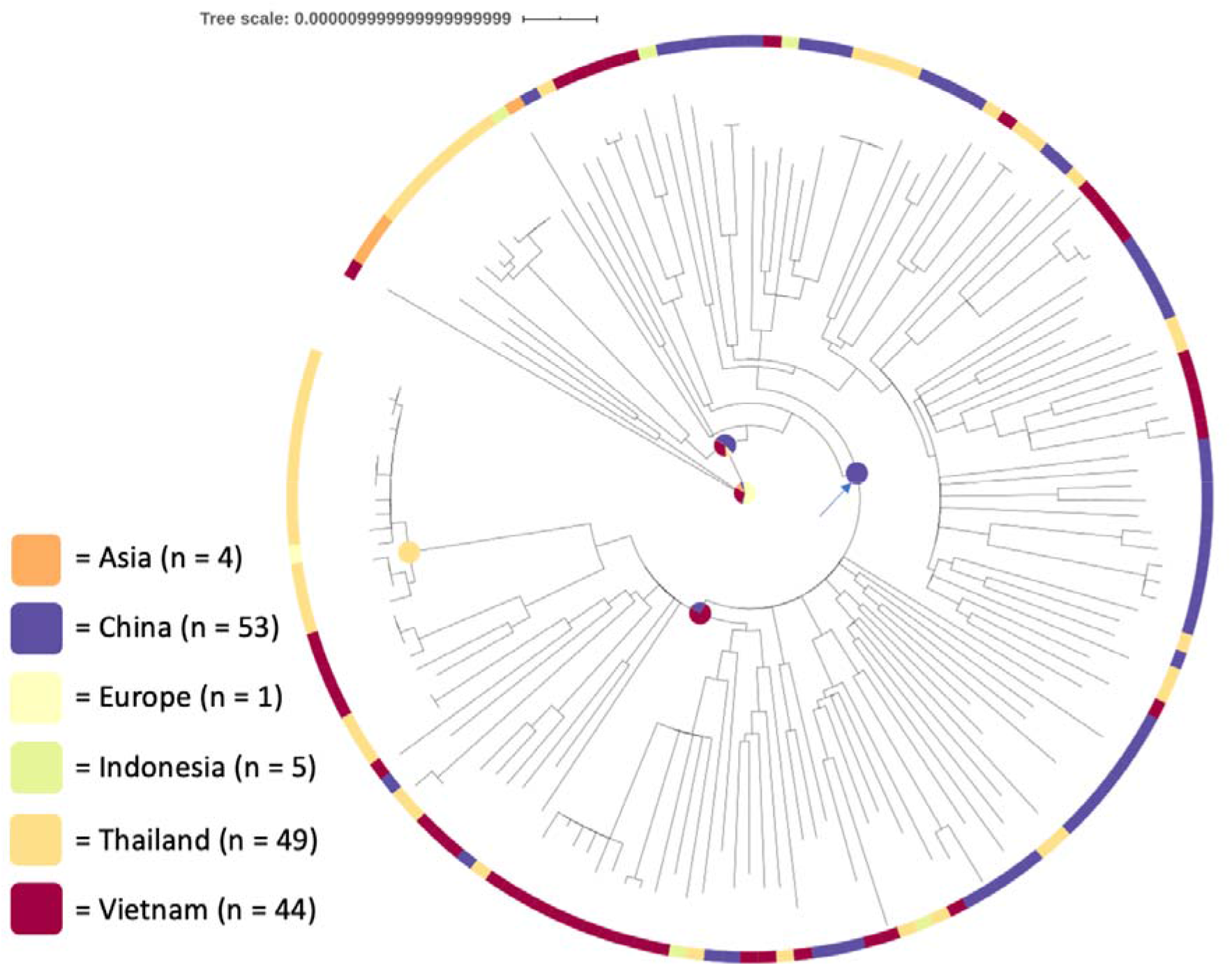
A maximum likelihood phylogeny of 152 L4.4.2 genomes. SIMMAP assignment of the MRCA of the entire sub-lineage is ambiguous, but there is a sub-clade of 135 genomes (indicated with an arrow) that was assigned to China with high confidence, and then a transmission from China to Vietnam, and subsequently from Vietnam to Thailand. Internal node annotations as per Figure 3.

#### L4.5

The third East/Southeast Asian sub-lineage was L4.5, with 270 genomes forming a sub-clade identified by TreeBreaker (Figure 5) (two basally branching genomes within this clade were assigned as L4 by tb-profiler). There were 101 Thai, 99 Chinese, 46 Vietnamese, 13 Indonesian, 6 other Asian and 5 European genomes in this sub-clade. The MRCA of this sub-clade was assigned to China (89%), while the parent of the MRCA was assigned to Europe (92%). According to the SIMMAP analysis, there were two high confidence transmissions from China into Thailand which resulted in two large Thai sub-clades of 26 and 13 genomes. There were 10 other China to Thailand transmissions which resulted in smaller sub-clades of 2 to 8 sampled genomes, containing a total of 34 Thai samples. In contrast, 37 of the 46 (80%) L4.5 sampled in Vietnam descend from a single introduction from China. In the sub-sampling analysis, 85.6% of replicates placed the MRCA in China, and 14.4% in the UK.

**Figure 5:**
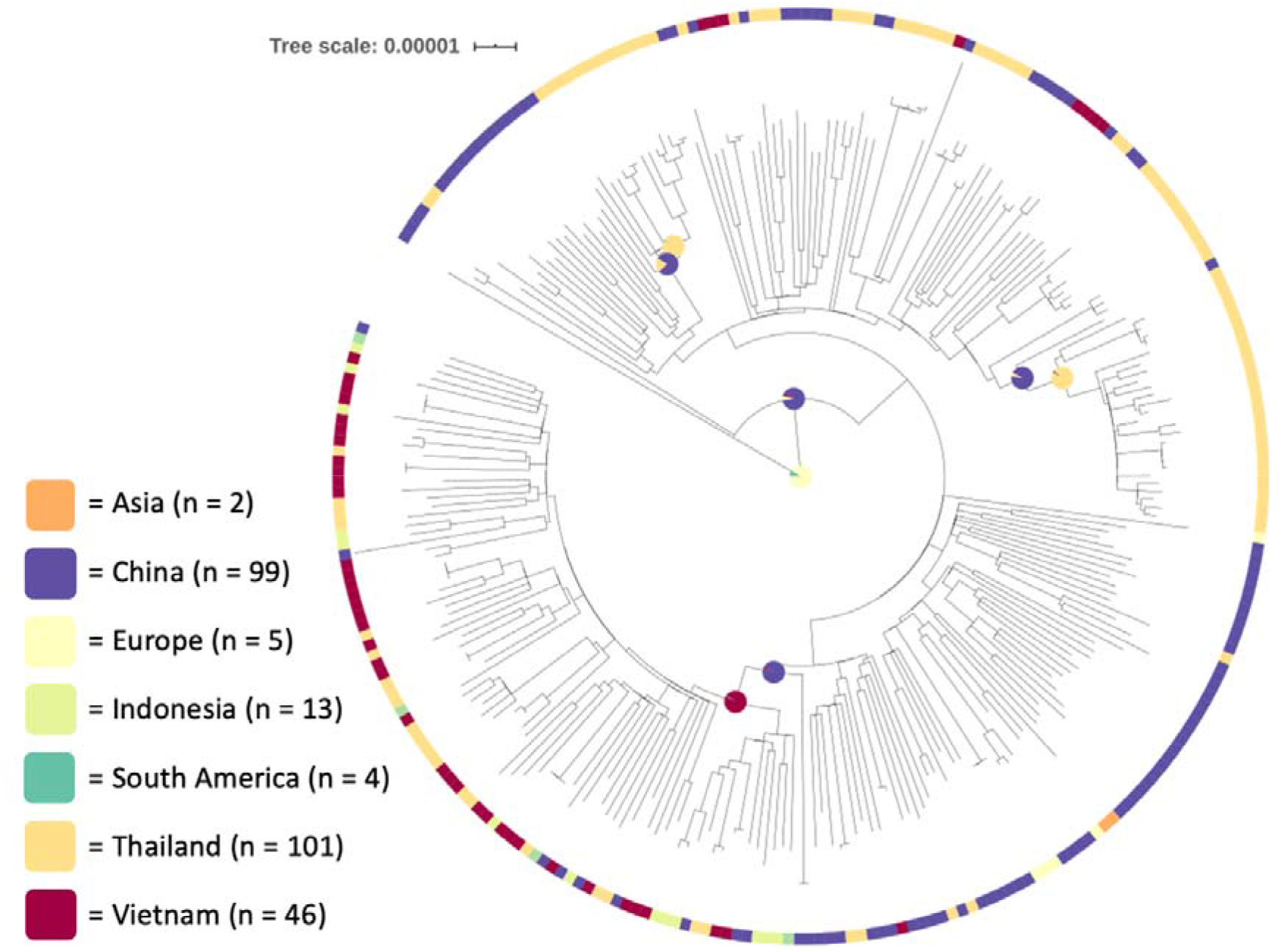
A maximum likelihood phylogeny of 270 L4.5 genomes. SIMMAP analysis indicates that there was a transmission from Europe to China. From China there were multiple transmissions to Thailand and separately to Vietnam. There were multiple other high confidence transmissions, enumerated in the manuscript text. Internal node annotations as per Figure 3.

### Global sub-clades

#### L4.1.2 & L4.1.2.1

Twenty five percent of Lineage 4.1.2 and its sub-lineage L4.1.2.1 were from Southeast Asia (Supplementary Table 2: A full line list of all lineage 4 readsets included in this analysis Supplementary Table 3, Figure 2). The MRCA of a sub-clade of 284 genomes identified by TreeBreaker was placed in Vietnam (64%) or Europe (21%) by SIMMAP, while the parent of the MRCA was placed in Europe (74%) or Vietnam (16%) (Supplementary Figure 2). SIMMAP confidently placed the MRCA of a sub-clade of 255 of the 284 genomes in Vietnam (96% confidence), but then a sub-clade of 234 of these 255 genomes had an MRCA of Europe (89% confidence). There were three sub-clades that were high confidence (>80%) transmissions from Europe to Vietnam, leading to a total of 17 sampled Vietnamese genomes. Of the 64 Indonesian L4.1.2.1 genomes, 37 were in a sub-clade of 64 genomes for which the MRCA was likely in Indonesia (70%) while the parent node of the MRCA was probably in Europe (71%).

#### L4.4.1.1

L4.4.1.1 is a global lineage, with 26% of genomes coming from Southeast Asia. There were two Southeast Asian clades with more than two isolates within L4.4.1.1; one sub-clade consisted of 5 Vietnamese isolates. SIMMAP identified a South American location for the parent node of this Vietnamese sub-clade, as it was nested within a sub-clade of Brazilian TB genomes (Supplementary Figure 3). The second Southeast Asian L4.4.1.1 sub-clade consisted of eight Indonesian isolates, for which the parent node of the MRCA was placed in North America by SIMMAP, as many of its close phylogenetic neighbours were isolated in Canada.

#### L4.4.1.2

L4.4.1.2 is a small clade that was 65% Southeast Asian, with 54% from Indonesia and 11% from Vietnam. While this lineage is well established in Indonesia, there was considerable uncertainty as to the location of the parent node of the MRCA, with SIMMAP analysis identifying South America (48%) or Europe (38%) being the most plausible locations of origin.

#### L4.8

L4.8 was 28% Southeast Asian, with Indonesia contributing 12%, Vietnam 11% and Thailand 4%. There was one sub-clade of 47 genomes for which SIMMAP identified the most likely location of the MRCA as Indonesia (76%), but the location of the parent of the MRCA was uncertain (Europe 42% or Indonesia 36%). There was another sub-clade with seven Indonesian genomes which represented a high confidence (99%) transmission from Europe to Indonesia. One Vietnamese sub-clade of seven genomes was the result of a transmission from Europe (99%), while another sub-clade of 4 genomes was from Indonesia (94%). There was a sub-clade of five Thai genomes which was also a direct transmission from Europe (100%).

#### L4.3 & L4.3.4.1 & L4.3.4.2

There are only 35 L4.3 genomes in our analysis, but 74% of them were Indonesian. While the MRCA of a sub-clade of 25 of them was confidently placed in Indonesia (97% confidence), the parent of the MRCA was either in South America (60%) or Europe (27%). L4.3.4.1 and L4.3.4.2 are associated with South America, with 56% and 57% of their genomes coming from that continent. Two sub-clades of Southeast Asian genomes were both from the Philippines; one of 11 Filipino genomes where the parent of the MRCA was placed in South America (99%) and a sub-clade of 17 genomes, of which 8 were Filipino, and where the MRCA was in the Philippines (100%) and the parent of the MRCA was either in South America (70%) or the Philippines (30%).

## Discussion

Our findings extend the idea of the “out of Europe” spread of *M. tuberculosis* L4 by showing that there were historical movements of *M. tuberculosis* L4 between countries in East and Southeast Asia. The sub-clades of *M. tuberculosis* L4 transmitted between East and Southeast Asian countries continue to be important contributors to the burden of *M. tuberculosis* L4 disease in the continent. While the limitations of the current sampling frame cause uncertainty around the precise order of which country “transmitted” to which, the data suggest that China was the intermediary between Europe and Southeast Asia for two of the Asian sub-clades, L4.2.2 and L4.5.

A major strength of our study is that we have carried out a combined analysis of *M. tuberculosis* L4 datasets from every Southeast and East Asian country for which they are available and placed them into a global context. Furthermore, we have employed a novel approach of using TreeBreaker as a screening tool to identify clades of interest for more in depth phylogeographic investigation in this large dataset. The main limitation of our analysis is that phylogeographical investigations are very susceptible to sampling bias so we should be cautious in our interpretation. This is especially true when analysing data that comes from studies with very different sampling frames, such as those analysed here. In the sub-sampling analysis, 14% of sub-samples placed the MRCA of L4.5 in the UK, reflecting the sensitivity of phylogeographic analysis to the small number of relatively deeply branching L4.5 genomes from the UK. The uncertainty in the location of the MRCA of L4.4.2 in the sub-sampling analysis was mirrored by the uncertainty in the SIMMAP assignment for this MRCA and suggests a complex migratory history that cannot be explained by the currently sampled genomes. Another limitation is that few countries in our analysis have nationally representative genome collections, with most represented by genomes from only one city or region.

Our study extends the findings of recent studies. Brynildsrud et al. and Holt et al. found that L4 has been introduced to Vietnam multiple times, with the first time being from Europe at the beginning of the 13^th^ Century (9,25). However, these analyses did not include genomes from Thailand, China, or Indonesia, and so their ability to identify intra-Asian transmissions was limited. Liu et al. analysed Chinese and Vietnamese L4, but their focus was on L4 within China, and they only noted that “closest branches to the strains sampled from Vietnam were mostly collected in South China” (15). An analysis of all lineages of *M. tuberculosis* in Africa and Eurasia found that Southeast Asia was the most connected region in terms of *M. tuberculosis* migrations globally [O’Neill 2019]. While O’Neill et al. found this to be primarily driven by Lineage 2, the high level of connectedness in Southeast Asia is also reflected in the dynamic picture of L4 transmission we have identified. The impact of historical population movements on the distribution of L4 has been well described (8,9,31). From our analysis, we can see the impact of pre-colonial, colonial and post-colonial relationships in the L4 phylogeny, with “transmissions” from South America to the Philippines, and between Indonesia and the Netherlands. The South America to the Philippines transmissions could have been true direct transmissions, mediated by the colonial trade links such as the Manilla galleons which sailed between the Philippines and Spanish colonies in central and South America, or could represent transmission from a common, unsampled source (i.e. historical Spain). The evidence of transmissions from Indonesia to the Netherlands is consistent with the historical Dutch colonisation of Indonesia, and demographic and cultural connections that persist to this day. L4.4.1.1 has been reported as transmitted from French-Canadian fur traders to Western Canadian First Nations people in the 18^th^ and 19^th^ centuries (45), and in Polynesia, linked with European whalers and other merchants (46). Here, we report that this lineage is a major cause of *M. tuberculosis* L4 cases in Indonesia. This adds to the remarkably diverse destinations of this well-travelled sub-lineage. While the SIMMAP analysis identified a North America to Indonesia transfer, based on the Canadian genomes sampled by Pepperell et al., a more historically congruent explanation could be speculated as both the Canadian and Indonesian sub-clades originating from a clonal population in France, with the possibility that the transfer to Indonesia could be connected to the French administration of Indonesia between 1806-1811 (47). As the French *M. tuberculosis* population underwent a major bottleneck in the 20^th^ century, it is unlikely that we will have strong phylogenetic evidence of seeding from France without historical French genomes.

One major implication of our findings, building on those of Liu et al. and Brynildsrud et al., is that multiple sub-lineages of L4 have been circulating in Asian populations for hundreds of years. Considering the hypothesis that *M. tuberculosis* is undergoing host population specific adaptation (48,49), in future work it would be interesting to look for signals of adaptation to that specific host population.

While the findings reported here enrich our understanding of L4 in Asia, having consistent sampling frames between different countries would increase the certainty of the conclusions we can draw. Therefore, in future research, carrying out structured surveys like those of Liu et al., or using unique isolate collections from those such as National TB Prevalence surveys, would provide a more comprehensive picture of TB in the region. In addition, the analysis of historical TB genomes has improved our understanding of the 19^th^ Century European TB epidemic. Having further contextual genomes from a broader swathe of 19^th^ Century Europe would provide very interesting context for these samples.

## Methods

### Data download

We carried out a literature review to identify papers that reported either *M. tuberculosis* genomes from Southeast Asia or globally representative *M. tuberculosis* L4 genomes. We used this search strategy to identify as many Southeast Asian L4 genomes as possible (by including all studies from the region), while also including representatives of global diversity.

### Data processing

Downloaded data were processed with bbduk v38,96 (50) in order to remove adapters and low quality sequencing regions ‘bbduk.sh ref=adapters.fa in=!{forward} in2=!{reverse} out=!{pair_id}_bbduk_1.fastq.gz out2=!{pair_id}_bbduk_2.fastq.gz ktrim=r k=23 mink=11hdist=1 tbo tpe qtrim=r trimq=20 minlength=50’. Trimmed fastqs were then analysed with tb-profiler (51) in order to identify the lineage of each readset. Trimmed fastqs were also mapped against the H37Rv reference genome (NCBI accession NC000962.3) using bwa mem v0.7.17-r1198-dirty (52). SNPs were called with GATK version 3.8-1-0-gf15c1c3ef in unified genotyper mode (53). Positions where the majority allele accounted for < 90% of reads mapped at that position, which had a genotype quality of <30, depth <5x, or mapping quality <30 were recorded as Ns in further analyses. A consensus genome was generated for each genome. These steps were carried out using the PHEnix pipeline (54).

### Phylogenetics and phylogeography

After consensus genomes were combined, we used snp-sites v2.5.1 to extract the variant positions, and then generated a neighbour joining tree of all samples with IQ-TREE v2.1.4-beta (55). The tb-profiler results were combined with the neighbour-joining tree and the lineage 4 genomes identified. A maximum likelihood phylogenetic tree of the L4 genomes was then derived using IQ-TREE with built-in model selection, and the inclusion of the number of invariant sites, as identified using snp-sites. TreeBreaker v1.1 (44) was used to identify internal nodes of the tree where there was a change in the distribution of phenotypes of interest at the tips that descended from that internal node. The phenotype of interest was the geographic location. To enable easy interpretation, separate TreeBreaker runs were carried out for Vietnam, Indonesia, China, Thailand, and all the preceding countries combined into a single category. The results from these different runs were combined for further analysis. SIMMAP analysis (56) was carried out using the make.simmap function from PhyTools (57) in the R statistical language (58) using RStudio (59). The fit of each model type (all rates different, symmetrical, and equal rates) was assessed using the fitMk function, and the model with the best fit used for the SIMMAP analysis. We ran 1000 simulations within SIMMAP. Nodes that were identified as being associated with changes by TreeBreaker were targeted for investigation in the output of SIMMAP. Trees were visualised with iTOL (60), and graphs drawn with ggplot2 (61).

### Sub-sampling

To investigate whether phylogeographical results identified in the full dataset were robust to differences in sampling, we sub-sampled 20 genomes from each of China, Thailand, Indonesia, and Vietnam, and all the European genomes available for each sub-clade, and carried out phylogeographical analysis using the TreeTime mugration command (62), and the location of the MRCA of the Asian sub-clade was extracted. This sub-sampling and phylogeographical analysis was repeated 1000 times.

### Conflicts of interest

The authors have no conflicts of interest to report.

## Acknowledgements

We would like to acknowledge Qingyun Liu and Carolien Ruesen for providing the year of isolation of their published genome sequences, and Prasit Palittapongarnpim and Lidya Chaidir for providing supplementary information on their *M. tuberculosis* genome sequences.

---

*Supplementary Table 2: A full line list of all lineage 4 readsets included in this analysis*

*Supplementary Table 3: Full lineage by country, the number of readset from each country belonging to each sub-lineage of Lineage 4*

**Supplementary Figure 1:**
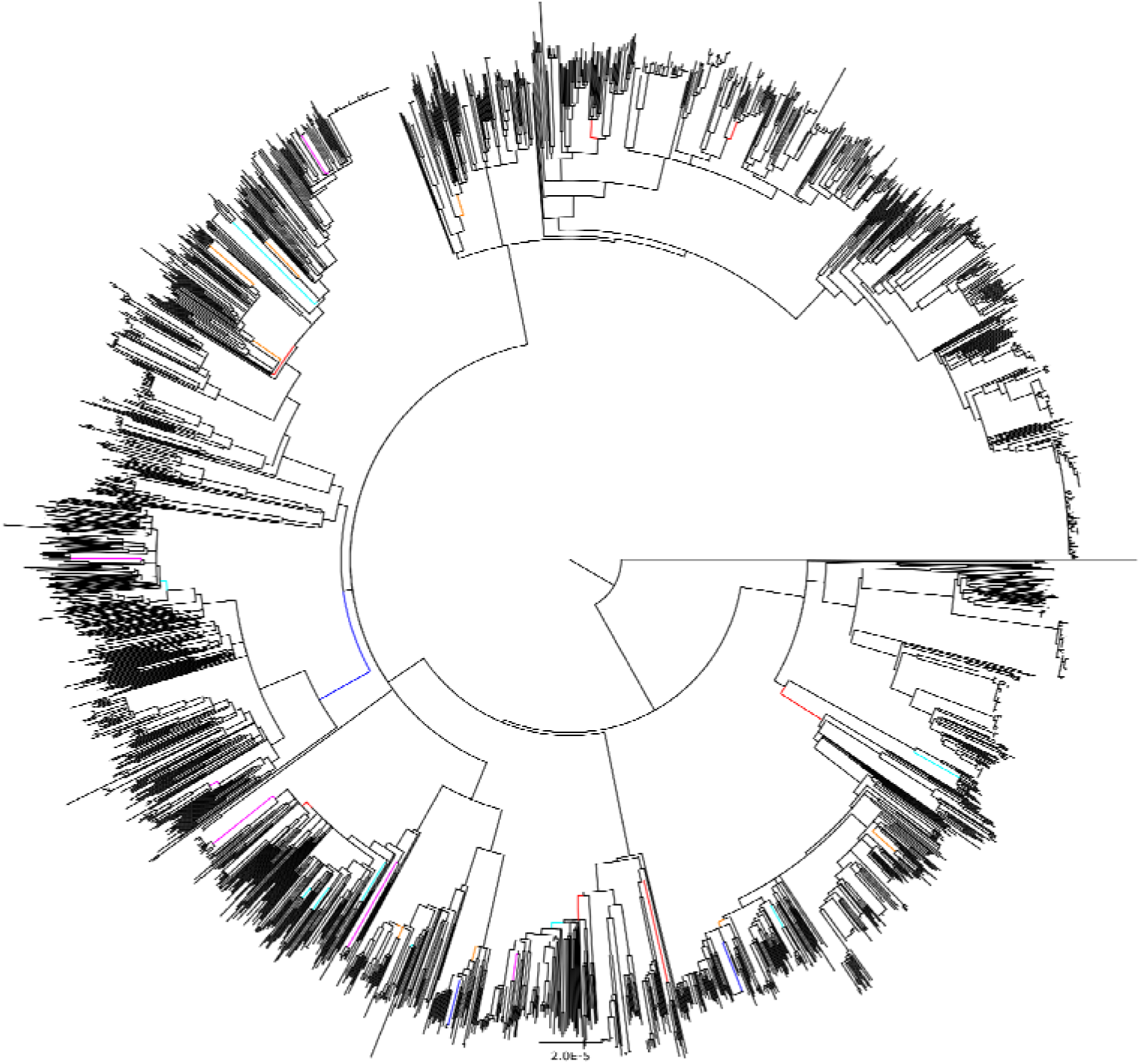
A maximum likelihood phylogeny of 2258 M. tuberculosis Lineage 4. Internal nodes are coloured when both TreeBreaker and SIMMAP analysis identified a change between geographic areas of interest on that branch. Colour coding is - red = a change to Southeast Asia & China, cyan = a change to Vietnam, orange = a change to Indonesia, magenta = a change to Thailand, royal blue = reversions from Asia to non-Asia.

**Supplementary Figure 2:**
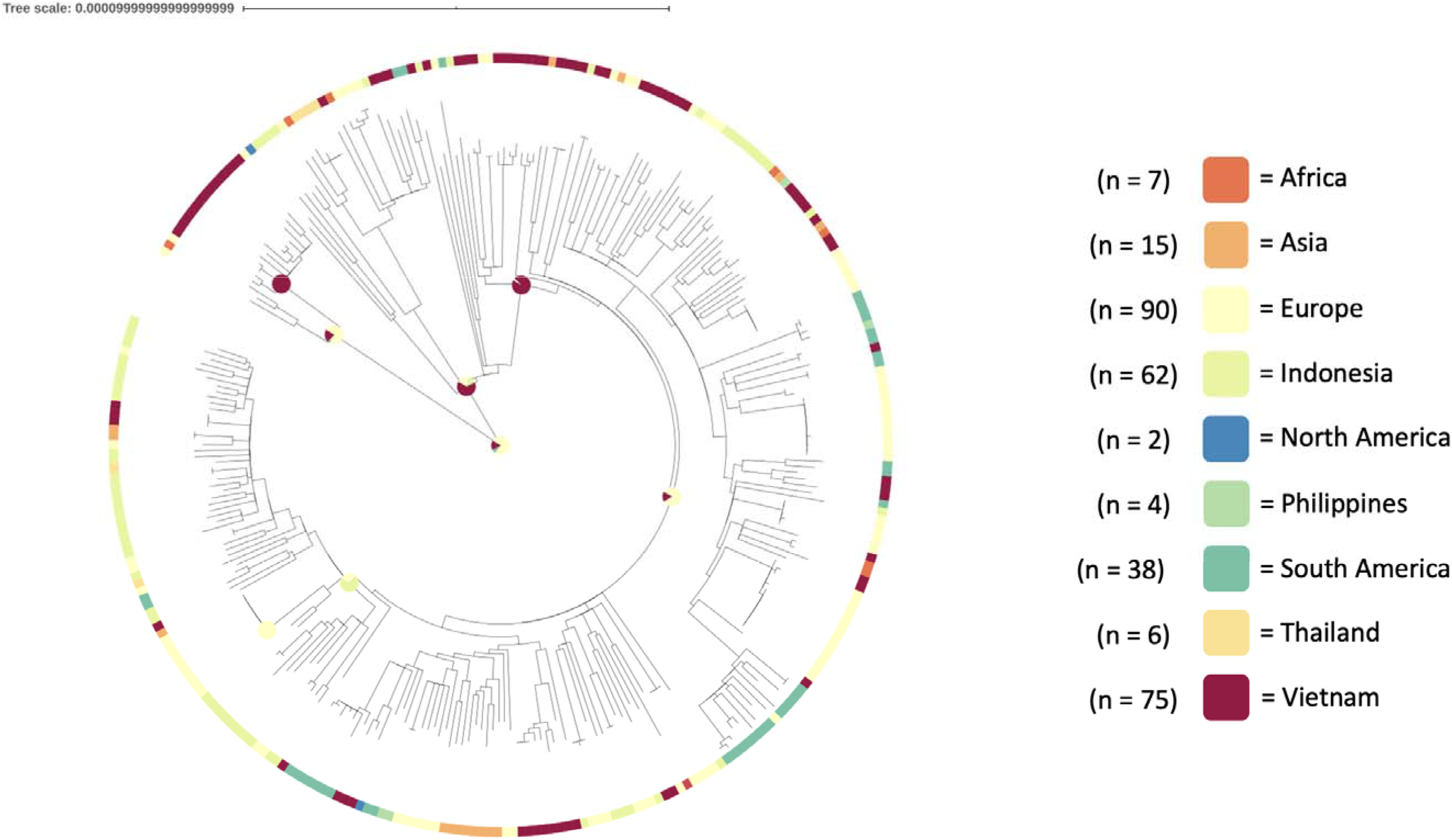
A maximum likelihood phylogeny of lineage 4.1.2 and descendants. Selected nodes are annotated with pie charts reflecting the placement of that node in SIMMAP analysis.

**Supplementary Figure 3:**
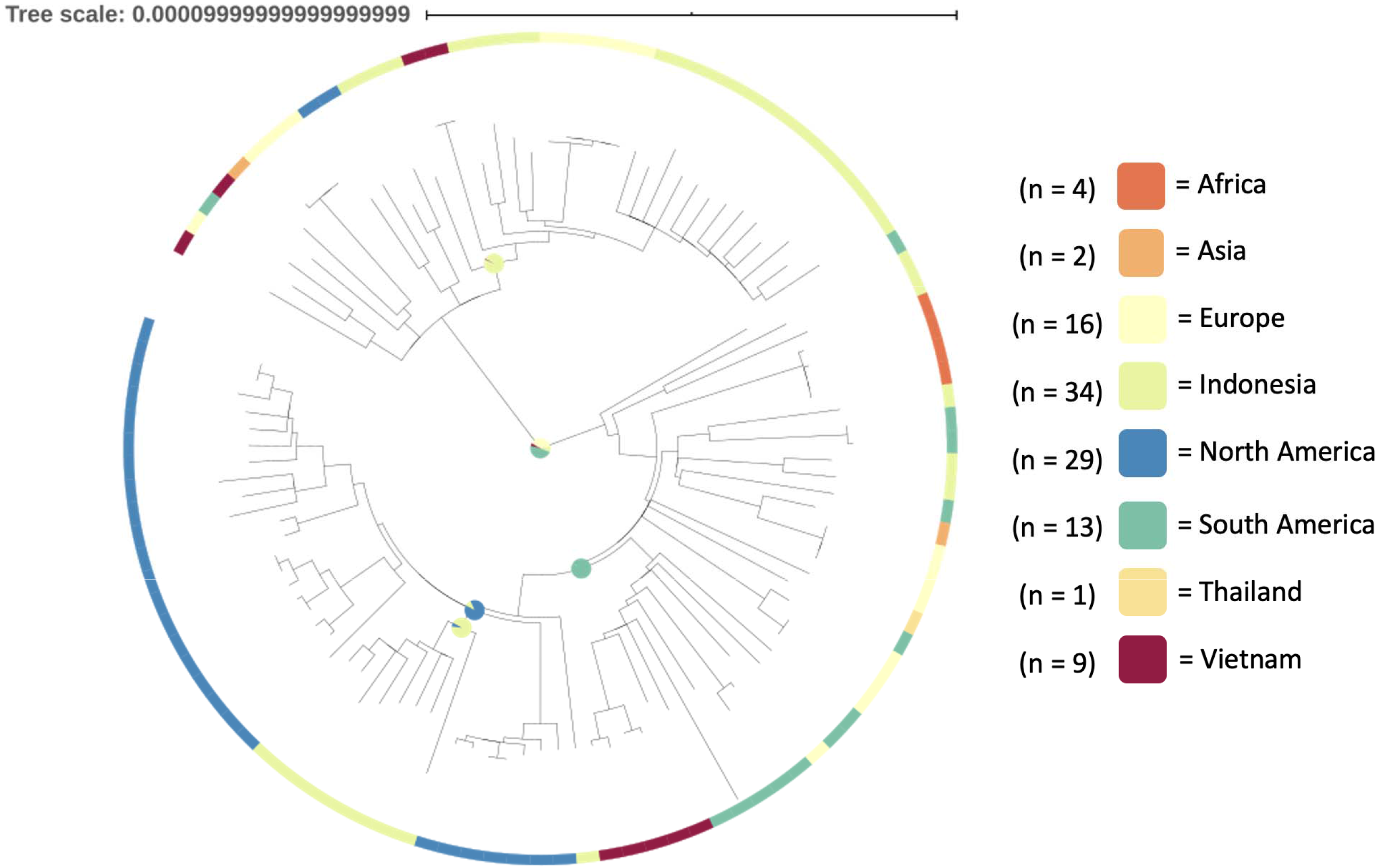
A maximum likelihood phylogeny of lineage 4.4.1.1 and descendants. Selected nodes are annotated with pie charts reflecting the placement of that node in SIMMAP analysis.

## Notes

### Competing Interest Statement

The authors have declared no competing interest.

